# CUADb-based oligonucleotide insecticides and RNA biocontrols: molecular bases and potential in plant protection

**DOI:** 10.1101/2024.03.13.584797

**Authors:** Vol Oberemok, Kate Laikova, Jamin Ali, Ilyas Chachoua, Nikita Gal’chinsky

**Affiliations:** Department of General Biology and Genetics, Institute of Biochemical Technologies, Ecology and Pharmacy, V.I. Vernadsky Crimean Federal University, Simferopol 295007, Republic of Crimea, Russia; (V.O.); (K.L.); College of Plant Protection, Jilin Agricultural University, Changchun 130118, China; (J.A.); Department of Molecular Biology and Genetics, Bilkent University, Ankara 06800, Turkey; (I.C.)

**Keywords:** oligonucleotide insecticides, CUAD biotechnology, DNA containment, RNA biocontrols, double-stranded RNA technology, RNA interference, next-generation insecticides, plant protection

## Abstract

Recent advances in molecular genetics, nucleic acid synthesis, and bioinformatics have provided novel opportunities for plant protection against insect pests. Currently, both DNA and RNA serve as active insecticidal ingredients, transcending their traditional role as carriers of genetic information. This novel activity is achieved through two fundamentally distinct mechanisms: DNA containment (DNAc), employing oligonucleotide insecticides based on contact unmodified antisense DNA biotechnology (CUADb), also known as ‘genetic zipper’ technology, and RNA interference (RNAi), employing RNA biocontrols based on double-stranded RNA (dsRNA) technology. The investigation of the molecular mechanism underlying the antisense activity of nucleic acids emerged in the early 1960s. While the antisense function of RNA in gene silencing through interference (RNAi) has been documented in the late1990s as an antiviral immune response in nematodes, the CUADb antisense approach initially emerged as a powerful strategy for pest control against lepidopterans in 2008. CUADb approach relies on disrupting rRNA biogenesis and ribosome production, a process entirely distinct from RNAi. The efficacy of these approaches appears to be species dependent: while CUADb demonstrates optimal activity against Sternorrhyncha (e.g., aphids, mealybugs, psyllids, scale insects), thrips, and mites, RNAi strategy shows a strong insecticidal potential against beetles from the Tenebrionidae and Chrysomelidae families. Here, we will review the differences between the two technologies, their mechanism of action and the current challenges facing their adoption.

## 1. Introduction

The relentless innovation in insect pest control drives continuous replacement of older chemistries (carbamates, organophosphates) by contemporary agents (neonicotinoids, diamides) (van den Berg et al. 2021; Sparks et al. 2021; Zhang et al. 2023). Yet the intractable challenge of genetic resistance ensures this cycle persists (Sparks et al. 2021; Mogilicherla et al. 2023), demanding insecticides with extended utility, heightened selectivity, enhanced biodegradability, and reduced carbon footprint. Biomolecules have recently emerged as promising tools that demonstrate high potential in overcoming these challenges (Christiaens et al. 2020; Oberemok et al. 2022; Palli 2023; Gal’chinsky et al. 2024). Both DNA and RNA have shown remarkable insecticidal effects, enabled by advances in molecular genetics, synthesis, and bioinformatics (Oberemok et al. 2018a; Rank and Koch 2021; Oberemok et al. 2023; Koeppe et al. 2023). This led to the development of two distinct technologies: oligonucleotide insecticides based on Contact Unmodified Antisense DNA Biotechnology (CUADb), also known as ‘genetic zipper’ technology, and RNA biocontrols utilizing double-stranded RNA (dsRNA) technology (Nitnavare et al. 2021; Gal’chinsky et al. 2024). Alongside CRISPR/Cas, all together they represent three core antisense approaches for insect control, each employing unique molecular mechanisms: RNAi functions through guide RNA-mRNA duplexes cleaved by Argonaute nuclease; CUADb operates via guide DNA-rRNA duplexes processed by DNA-guided rRNase such as RNase H1; and CRISPR/Cas combines guide RNA-genomic DNA complexes with Cas nuclease (Kumar et al. 2025). While all three exhibit taxon-specific efficacy enabling combinatorial strategies, their divergent molecular targets preclude universal application, increasing the chance of efficiently targeting different organisms.

After two decades of development, RNAi still lacks a standardized design algorithm for reliable insecticide development. Conversely, CUADb possesses a validated design framework (Oberemok et al. 2024a) with potential efficiency against up to 50% of all insect pests, demonstrating particular efficacy against Sternorrhyncha (e.g., aphids, mealybugs) and superorder Paraneoptera in general (Oberemok et al. 2025a), whereas RNA biocontrols show strongest activity mainly against Coleoptera. These independent ‘fraternal twins’, dsRNA technology and ‘genetic zipper’ technology (CUADb), hold synergistic potential for developing highly selective, next-generation pest control agents. This review addresses the paradigm shift toward nucleic acid-based solutions, with dedicated focus on CUADb as an innovative technology characterized by its novel DNAc mechanism and advanced insecticidal profile.

## 2. CUADb-based oligonucleotide insecticides

In 2008, our research group pioneered the use of short unmodified antisense DNA oligonucleotides as contact insecticides (Oberemok 2008). Initial work on the spongy moth (*Lymantria dispar*) demonstrated that effective gene silencing using oligonucleotide insecticides (oligoRING and oligoRIBO-11) depended critically on target gene and its expression levels (Oberemok et al. 2017, 2019). Subsequent research identified rRNA as a prime target, leading to the development of oligonucleotide insecticides based on 11-mer DNA sequences complementary to pest rRNA in 2019 (Kumar et al. 2025). rRNA is an optimal target due to its abundance (constituting ∼80-85% of cellular RNA) and the substantial energy investment (>60% of cellular energy) required for ribosome production and maintenance (Szaflarski et al. 2022; Deng et al. 2022). Insect rRNA comprises 28S rRNA (∼3900 nt), 18S rRNA (∼1920 nt), 5.8S rRNA (∼160 nt), 5S rRNA (∼120 nt), mitochondrial rRNAs including 16S rRNA (∼1140 nt) and 12S rRNA (∼600 nt) (Deng et al. 2022). The first successful rRNA targeting with unmodified antisense DNA oligonucleotides was achieved against 5.8S rRNA of *L. dispar* (Oberemok et al. 2019). CUADb-based oligonucleotide insecticides are designed using the DNAInsector program (dnainsector.com) or manually via GenBank sequences of pest pre-rRNA and mature rRNA. Synthesis employs the phosphoramidite method through liquid-phase or solid-phase synthesis using instruments such as the ASM-800 (BIOSSET, Russia), OligoPilot™ (Cytiva, Sweden), PolyGen 10-Column DNA Synthesizers, and others (Gal’chinsky et al. 2019).

Oligonucleotide insecticides demonstrate high efficacy against sternorrhynchans, thrips, and mites, with successful targeting documented across multiple species: 28S rRNA in *Unaspis euonymi, Dynaspidiotus britannicus, Icerya purchasi, Ceroplastes japonicus, Aonidia lauri*, and *Coccus hesperidum* (Gal’chinsky et al. 2020, 2023, 2024; Useinov et al., 2020; Oberemok et al., 2022, 2025a); 18S rRNA in *Pseudococcus viburni* (Novikov et al. 2023); mitochondrial 16S rRNA (Oberemok et al. 2025b); and ITS2 regions in *Macrosiphoniella sanborni, Schizolachnus pineti* (Puzanova et al. 2023; Oberemok et al. 2024b), *Trioza alacris* (Oberemok et al. 2024c); and also *Tetranychus urticae* (Gavrilova et al. 2025), demonstrating high insecticidal potential for oligonucleotide acaricides (Kumar et al. 2025). A single contact treatment at 100 ng/μl typically achieves approximately 80% mortality in pests within 3-14 days (Oberemok et al. 2023, 2024a). While highly effective against hemipterans and moderately effective against lepidopterans including *L. dispar*, efficacy is notably lower against coleopterans such as *Leptinotarsa decemlineata* and requires further investigation of their resistance to this approach (Oberemok et al., 2018b). The 11-mer length provides species specificity with a uniqueness frequency exceeding 1/4.19·10 ^m^, covering most agricultural applications (Oberemok et al. 2022). rRNA’s dual advantages of abundance and inter-species variability make it superior to less concentrated cellular mRNAs as a target. Contact delivery (CUADs) outperforms oral delivery (ODUADs) due to digestive DNases and extra-oral salivary barriers in hemipterans (Cao et al. 2018).

The 2008 discovery of contact oligonucleotide insecticides was unexpected, contradicting established views that unmodified oligodeoxyribonucleotides are rapidly degraded by DNases (Dias and Stein 2002) and that rRNA resists RNase H-dependent antisense DNA degradation (Will and Lührmann 2001; Bachellerie et al. 2002). Previous research focused exclusively on gene downregulation, overlooking insecticidal potential (Dias and Stein 2002; Oberemok 2008). Oligonucleotide insecticides (briefly olinscides or DNA insecticides) refuted these assumptions through a two-step DNA containment (DNAc) mechanism (Figure 1, left part): initial target rRNA ‘arrest’ halts ribosome function and triggers rDNA transcription hypercompensation, followed by degradation of target rRNA by DNA-guided rRNases such as RNase H1 (Kumar et al. 2025; Gal’chinsky et al. 2024). This ‘genetic zipper’ mechanism (Oberemok et al. 2024c) induces metabolic shifts toward lipid-based energy synthesis, enhancing biogenesis of ribosomes and ATP production. Ultimately, widespread kinase downregulation – including mTOR, which regulates ribosome biogenesis through mTORC1 – causes ‘kinase disaster’ due to ATP insufficiency, while significant RNase H1 upregulation occurs throughout DNAc (Oberemok et al. 2025a).

**Figure 1.**
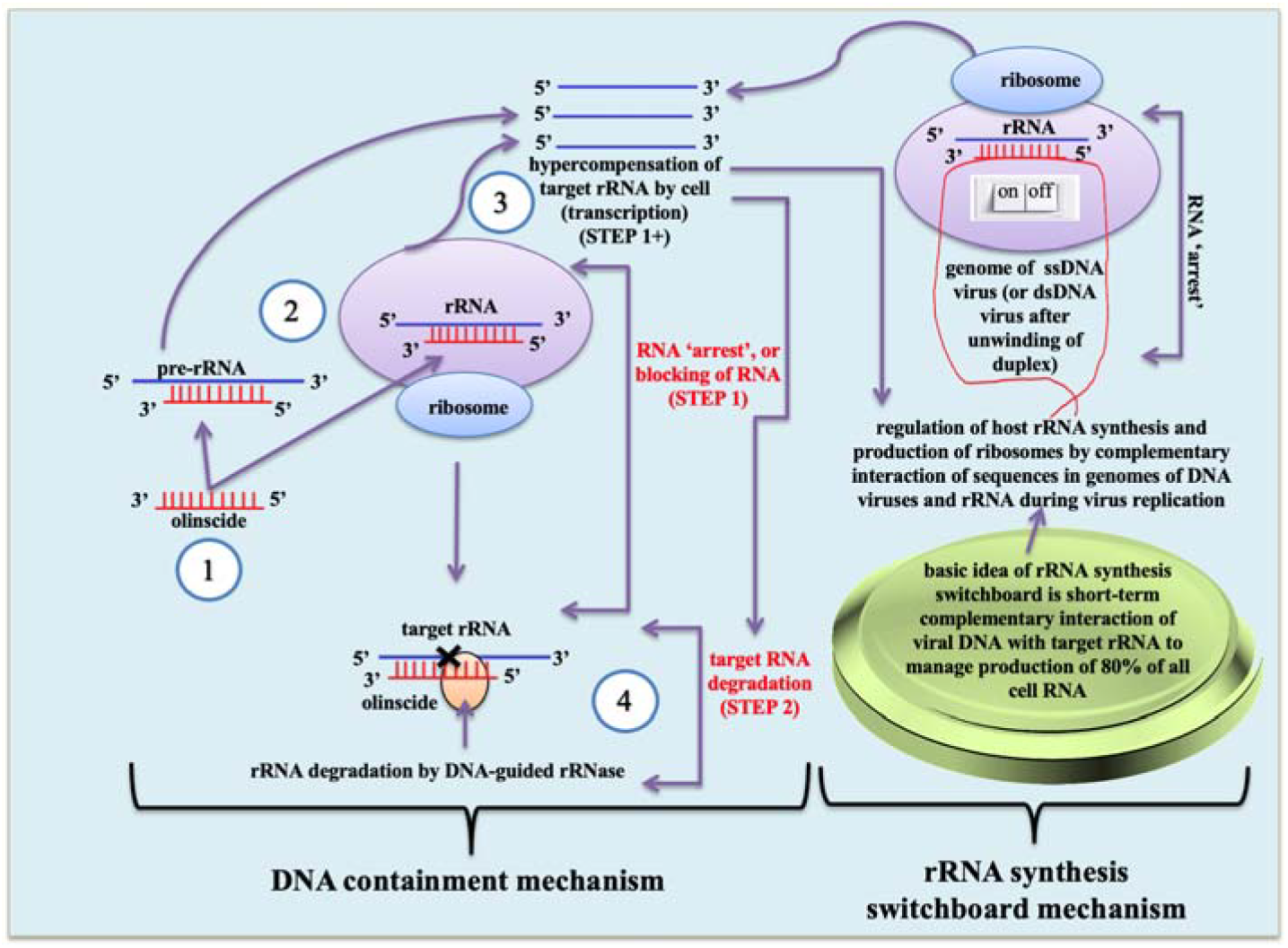
Mechanisms underlying oligonucleotide insecticides and rRNA regulation. Left: DNA containment mechanism (DNAc) for insect pest control. (1) Antisense DNA oligonucleotides complementary to pest mature rRNA or pre-rRNA are applied as insecticides. (2) Duplex formation occurs between the oligonucleotide and target rRNA/pre-rRNA. (3) In the first step of DNAc, antisense DNA ‘arrests’ rRNA or pre-rRNA, leading to hypercompensation and containment of ribosome biogenesis (‘arrested ribosomes’). (4) In the second step, DNA-guided rRNase cleaves the target rRNA/pre-rRNA, reducing its concentration. Right: rRNA synthesis switchboard mechanism, where DNA viruses regulate host rRNA synthesis and ribosome production by introducing complementary sequences of viral DNA, influencing up to 80% of rRNA output.

In 2011, three years after this discovery, Wang et al. (2011) adapted the contact concept for dsRNA insecticides. Crucially, DNAc downregulates key RNAi enzymes (DICER1, Argonaute 2, DROSHA), underscoring its mechanistic distinction from RNAi (Oberemok et al., 2025a). Thus, ssDNA in CUADb and dsRNA in RNAi represent complementary but fundamentally distinct antisense technologies with synergistic potential in pest management (Kumar et al. 2025) (Table 1).

**Table 1.** Comparative overview of CUAD biotechnology and dsRNA technology, highlighting differences in effectors, mechanisms, synthesis methods, and pest targets.

| Key features | CUAD biotechnology (‘genetic zipper’ technology) | dsRNA technology |
| --- | --- | --- |
| Effector | ssDNA | dsRNA |
| Mechanism | DNA containment | RNA interference |
| Mode of action | ssDNA:rRNA | ssRNA:mRNA |
| Nuclease involved | DNA-guided rRNase (such as RNase H1) | Argonaute |
| Synthesis | Liquid-phase solid-phase oligonucleotide synthesis based on phosphoramidite chemistry | Large-scale cell-free biomanufacturing (cell-free dsRNA production) |
| Target pests with best outcome | Sternorrhynchs | Beetles |

A working hypothesis proposes that DNA and RNA viruses may exploit the rRNA gene expression hypercompensation observed during the initial step of the DNA containment mechanism to increase cellular ribosome numbers essential for viral replication (Oberemok et al. 2025a). Bioinformatics studies indicate viral hijacking of host genes is common (Cerio et al. 2020). Given rRNA’s dominance (∼80% of cellular RNA) and the significant energy investment (>60%) in ribosome production (Deng et al. 2022; Szaflarski et al. 2022), DNA viruses likely co-opted rRNA-like sequences during co-evolution (Evseev et al. 2021) but their function is still unknown.

Most DNA viruses replicate within the nucleus and interact with the nucleolus – a site critical for ribosome assembly and virus life cycle (Salvetti et al. 2024). Nucleoli facilitate rRNA synthesis, processing, and pre-ribosomal subunit assembly (Rawlinson et al. 2018). Our studies suggest that specific DNA virus genome fragments could regulate host rRNA synthesis via an ON/OFF switch mechanism, using complementary interactions between viral DNA and host rRNA to initiate ribosome biogenesis with minimal energy expenditure (Oberemok et al. 2025a). This adaptation is likely to be very important since helps quickly and efficiently bypass host limitations. For example, viruses may need to replicate while the host cell is in resting phase (e.g., G2/M resting phase), where polymerases may not be readily available. Also some large DNA viruses replicate in the cytoplasm, a region of the cell that lacks the necessary host polymerases for DNA replication, which is typically carried out in the nucleus. Thus, fast and elegant rRNA synthesis switchboard mechanism for turning on the ribosome biogenesis is required for DNA viruses in both nucleus and cytoplasm (Figure 1, right part). Additionally, DNAc may function within the innate immune system against ssDNA viruses (their major vectors are hemipterans) and DNA viruses infecting hemipterans (Barnes and Turner 2016; Wang and Blanck 2021; Guo et al. 2022). DNAc can help degrading virus mRNAs using DNA fragments of partially cleaved virus genome and enhancing ribosome biogenesis to counteract viral pathogen (Gal’chinsky et al. 2024). Of note, in our investigations all DNA oligos with either high or moderate complementarity to target rRNA initiated rRNA hypercompensation, but subsequent substantial rRNA degradation and insect mortality occurred only when the oligo sequence perfectly matched the rRNA. Also long DNA oligos with either high or moderate complementarity triggered stronger rRNA hypercompensation than short ones (Oberemok et al., 2025a). Thus, viruses obviously will tend to undergo microevolution toward having long but somewhat similar complementary sequences to host rRNAs rather than highly similar complementary sequences. This strategy will likely trigger robust rRNA biogenesis without risk of its further substantial degradation and death of the host. Interestingly, we found that many RNA viruses found in GenBank, including those of hemipterans, also contain long (>100 nt) rRNA-like sequences of insect pests located either near or in the locus of RNA-dependent RNA polymerase gene performing virus replication function. Obtained data indicates potential role of these sequences in regulation of rRNA synthesis and ribosome biogenesis required for fast synthesis of RNA-dependent RNA polymerase and subsequent replication of RNA viruses and managed through a process similar to rRNA synthesis switchboard mechanism (Figure 1, right part) via viral RNA-host rRNA interactions.

From practical point of view, targeting conserved regions of insect pest rRNA genes for oligonucleotide insecticide design delays target-site resistance development, as mutations occur less frequently in these areas (Sharma et al. 2014; Oberemok et al. 2019). When resistance develops, new effective CUADb insecticides could be designed by shifting the target site adjacent to the resistant region within the rRNA genes (Gal’chinsky et al. 2024). Notably, current CUADb-based pesticides demonstrate efficacy against approximately 30% of the most resistant arthropods (Whalon and Mota-Sanchez 2015), showing substantial activity against Sternorrhyncha and spider mites (Kumar et al. 2025).

Oligonucleotide insecticides undergo rapid and clean biodegradation in the environment (soils, water, plants) via abiotic factors (temperature, pH, salinity, UV radiation) and biotic factors (microbes, extracellular enzymes) (Barnes and Turner 2016; Oberemok et al. 2024d). Deoxyribonucleases in tissue homogenates of *Lymantria dispar, Leptinotarsa decemlineata, Icerya purchasi, Dynaspidiotus britannicus, Aonidia lauri*, and their host plants (*Quercus pubescens, Solanum tuberosum, Pittosporum tobira, Laurus nobilis*) degrade olinscides within 24 hours at 27°C (Oberemok et al. 2018b, 2019; Gal’chinsky et al., 2023, 2024), while *Macrosiphoniella sanborni* nucleases achieve degradation within 1 hour (Puzanova et al., 2023). Minimal negative impacts are observed on non-target organisms including *Triticum aestivum* (Oberemok et al. 2013; Nyadar et al. 2018), *Quercus robur, Malus domestica* (Zaitsev et al. 2015), *Manduca sexta, Agrotis ipsilon* (Oberemok et al. 2015), and *Galleria mellonella* (Oberemok et al. 2019). Unprecedented selectivity is evidenced by dramatic efficacy loss following single nucleotide substitutions (Puzanova et al. 2023; Gal’chinsky et al. 2024), although tolerance for non-canonical base pairs (A:C, C:A, G:U) must be considered during design (Gal’chinsky et al. 2024; Oberemok et al. 2024b). Efficacy extends to mixed pest populations (Gal’chinsky et al. 2024), and modern liquid-phase synthesis minimizes greenhouse gas emissions (nitrogen oxide, methane, carbon dioxide) compared to solid-phase methods (Gal’chinsky et al. 2023), addressing climate concerns.

In conclusion, overcoming widespread pesticide resistance requires the development of novel and highly specific pesticide classes to reduce environmental impact. Oligonucleotide insecticides represent a remarkable advancement using unmodified antisense oligonucleotides. Mirroring the trajectory of therapeutic oligonu-cleotides – which overcame early challenges through sustained research and investment (TriLink BioTechnologies 2025) – CUADb-based pesticides offer selective action, rapid biodegradation, and increasingly cost-effective production, presenting a highly promising approach (Kumar et al. 2022). Based on our estimations, the ‘genetic zipper’ technology (CUADb) can potentially control up to 50% of all insect pests using a simple and flexible algorithm (Oberemok et al. 2024c). Formulation additives (spreaders, adhesives, penetrators, UV protectants) may enhance efficacy for other insect orders, pending environmental safety assessment. Future research will undoubtedly uncover deeper mechanistic details of DNAc and its potential. CUADb paves the way for sustainable, xenobiotic-free agriculture, positioning oligonucleotide pesticides as prospective cornerstone agents in pest control (Oberemok et al. 2025b).

## 3. RNA biocontrols

The foundation of RNA interference (RNAi) was established by Andrew Fire and Craig Mello’s seminal discovery that antisense strand within double-stranded RNA (dsRNA) mediates sequence-specific gene silencing – a breakthrough recognized with the 2006 Nobel Prize in Physiology or Medicine (Fire et al. 1998). This pivotal work catalyzed extensive biotechnological applications in agricultural and forestry research for plant protection. RNAi operates through post-transcriptional and translational suppression of gene expression, utilizing dsRNA molecules typically exceeding 200 base pairs in length that are complementary to target genes (Tomoyasu 2008; Svoboda 2020). The mechanism initiates when the RNase III enzyme (Dicer) cleaves long dsRNA into 20–30 nucleotide small interfering RNAs (siRNAs) (Carmell et al. 2002; Obbard et al. 2009; Koeppe et al. 2023). These siRNAs subsequently associate with Argonaute-family proteins to form RNA-induced silencing complexes (RISC) for cytoplasmic mRNA degradation, or RNA-induced transcriptional silencing complexes (RITS) for nuclear gene suppression (Wu et al. 2020; Müller et al. 2020). A parallel pathway involves PIWI-interacting RNAs (piRNAs; 23–36 nucleotides), germline-specific molecules that complex with PIWI-subfamily proteins to repress transposable elements through distinct silencing mechanisms (Bodemann et al. 2012; Chattopadhyay et al. 2022). Collectively, these RNAi pathways play crucial regulatory roles during insect ontogenesis (Figure 2), ultimately disrupting biosynthesis of specific proteins through targeted mRNA degradation or translational blockade.

**Figure 2.**
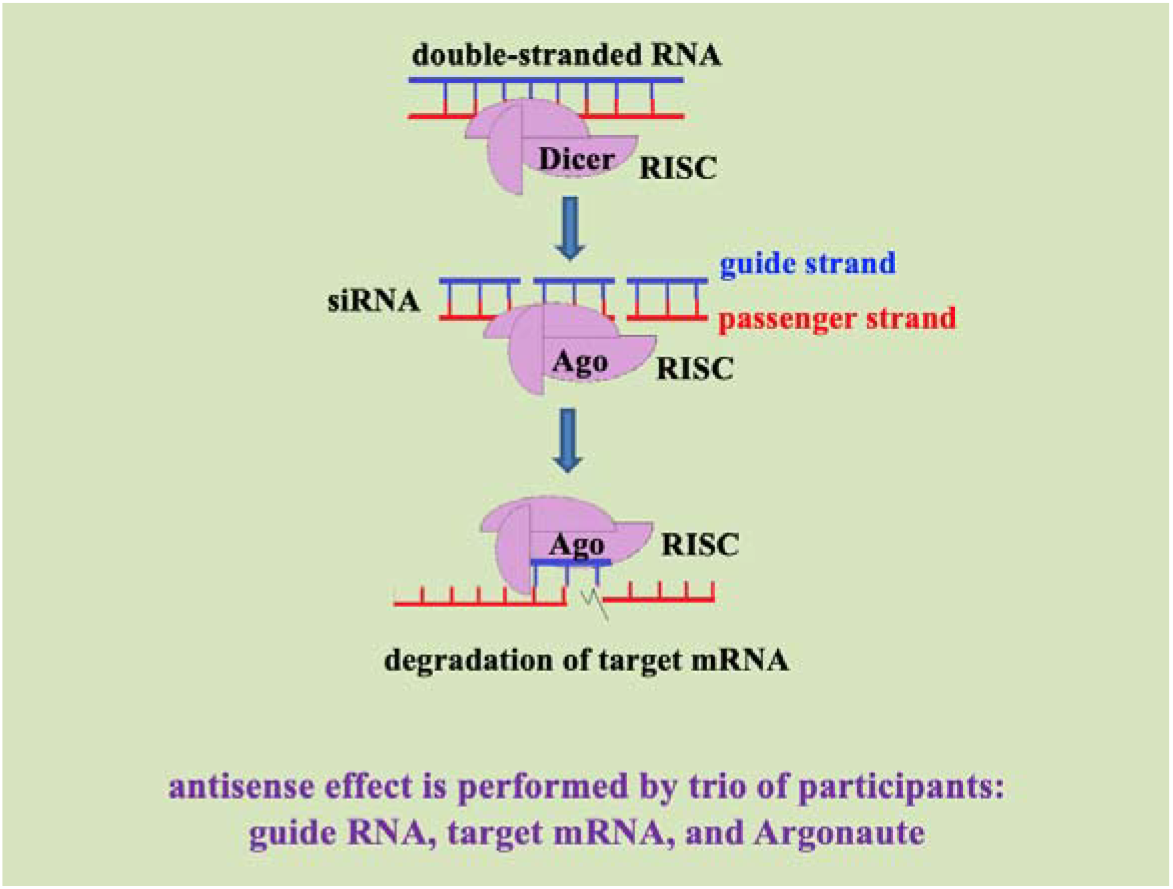
Main pathway of RNA interference (RNAi) utilized for the development of RNA-based biocontrols. Double-stranded RNA (dsRNA) is processed by Dicer into siRNAs, which are incorporated into the RNA-induced silencing complex (RISC). The guide strand directs Argonaute (Ago) to complementary target mRNA, resulting in mRNA cleavage and degradation. This mechanism, distinct from DNA containment (DNAc) and contact unmodified antisense DNA biotechnology (CUADb), forms the basis for RNAi-mediated pest control strategies.

RNAi efficacy exhibits profound taxonomic variation among economically significant insect pests. Coleopterans demonstrate consistently high systemic efficiency (Bodemann et al. 2012; Powell et al. 2017; Rodrigues et al. 2017), while Orthoptera (e.g., locusts) and Blattodea (cockroaches) generally exhibit robust responses (Irles et al. 2013; Santos et al. 2014; Li et al. 2021; Hoang et al. 2022). Efficiency is variable in Hymenoptera and Hemiptera due to limited systemic spreading in piercing-sucking insects, attributed primarily to deficient RNA-directed RNA polymerase (RdRP) activity (Christiaens et al. 2014; Jain et al. 2020, 2021). Lepidopterans and dipterans typically display low susceptibility to RNAi (Miller et al. 2008; Terenius et al. 2011). Consequently, commercial RNAi biopesticide development remains largely confined to coleopteran targets (Bramlett et al. 2020; Rodrigues et al. 2021; Koo et al. 2024), with target mRNAs spanning genes governing development, detoxification, and reproduction (Liu et al. 2020) (Table 1).

So far, two primary delivery strategies have been developed for RNAi: genetically modified (GM) plants expressing dsRNA, and topical application of formulated sprayable products (Ivashuta et al. 2015). GM approaches undergo protracted regulatory evaluation under biotechnology frameworks, whereas sprayable formulations – classified as biochemical pesticides – offer potential time and cost advantages, with registration timelines dependent on final product composition (Dietz-Pfeilstetter et al. 2021; Galli et al. 2024). Currently, few RNAi products are commercially deployed. The landmark SmartStax® PRO corn (US EPA-registered 2017) combines two Bt insecticidal proteins (Cry3Bb1 and Cry34/35Ab1) with DvSnf7 dsRNA targeting vesicle transport in corn rootworm (Head et al. 2017; EPA 2024). This triple-mode-of-action pyramid achieves 99% suppression of western (*Diabrotica virgifera virgifera*) and northern (*D. barberi*) corn rootworm larvae, though dsRNA-induced mortality manifests slower than Bt toxicity (≥5 days post-ingestion) (Head et al. 2017). The targeted Snf7 protein functions within the ESCRT-III pathway governing transmembrane protein sorting and degradation (Vaccari et al. 2009; Bolognesi et al. 2012), with documented absence of cross-resistance between DvSnf7 RNAi and Bt toxins (Moar et al. 2017; Khajuria et al. 2018; Reinders et al. 2023). Experimental GM crops demonstrate efficacy against diverse pests including fall armyworm (*Spodoptera frugiperda*; Xiong et al. 2013), aphids (Mao et al. 2014), Colorado potato beetle (*Leptinotarsa decemlineata*; Guo et al. 2018), and spider mites (*Tetranychus urticae*; Shen et al. 2017).

The first exogenous sprayable dsRNA biopesticide, Calantha™ (ledprona; IRAC Group 35), was commercialized by GreenLight Biosciences in 2023 against Colorado potato beetle (Pallis et al. 2023; GreenLight Biosciences 2025; Narva et al. 2025). Applied at 9.4 g/ha, it induces near-complete feeding cessation within 2-3 days and achieves 90-95% mortality in 1^st^-2^nd^ instar larvae within 11-26 days. Operational constraints include strict larval-stage specificity, necessity for sequential applications, and recommended rotation with insecticides possessing alternate modes of action. Proof-of-concept successes encompass numerous taxa: brown planthopper (*Nilaparvata lugens*; Zha et al. 2011), African sweet potato weevil (*Cylas puncticollis*; Prentice et al. 2017), tomato pinworm (*Tuta absoluta*; Camargo et al. 2016), oriental fruit fly (*Bactrocera dorsalis*; Sun et al. 2023), cotton mealybug (*Phenacoccus solenopsis*; Arya et al. 2021), diamondback moth (*Plutella xylostella*; Chen et al. 2021), fall armyworm (*Spodoptera frugiperda*; Li et al. 2024; Bera et al. 2025), white-backed planthopper (*Sogatella furcifera*; Mao et al. 2014; Guo et al. 2025), desert locust (*Schistocerca gregaria*; Schellens et al. 2022), and potato psyllid (*Bactericera cockerelli*; Lu et al. 2024) – though none have advanced beyond experimental stages.

Despite promising efficacy dsRNA technology is in its infancy and require broader adoption. There are main challenges restricting this adoption. Target gene selection lacks consensus due to compensatory cellular responses including protein stability, functional redundancy among paralogs, and transcriptional feedback loops (Ulrich et al., 2015; Buer et al., 2024; Cedden et al., 2024). Efficacy varies substantially with developmental stage, tissue type, and delivery method. The Spray-Induced Gene Silencing (SIGS) paradigm suffers mechanistic ambiguity – whether dsRNA uptake occurs primarily via integument penetration or oral ingestion and remains unresolved, confounding optimization efforts (Dalakouras et al. 2021). Off-target effects may arise from as few as 15 contiguous base matches (Powell et al. 2017; Chen et al. 2021) or sequence-independent immune activation (Hirai et al. 2004; Brutscher et al. 2017), compounded by formulants (nanocarriers, adjuvants) potentially enhancing non-target exposure (Castellanos et al. 2019; Reinders et al. 2023). Rapid environmental degradation occurs via microbial nucleases in soil/phyllosphere, UV radiation, and precipitation runoff (Dubelman et al. 2014; Parker et al. 2019), with insect-derived nucleases in hemolymph (e.g., *Spodoptera litura, Manduca sexta*) and saliva (*Acyrthosiphon pisum*) degrading dsRNA within 0.3–3 hours and interferes with robust insecticidal effect (Christiaens et al. 2014; Garbutt et al. 2013; Peng et al. 2018). While dsRNA production costs have decreased through cell-free platforms like GreenLight’s enzymatic NMP→NTP→dsRNA pipeline (Cunningham et al. 2020; Taning et al. 2020), it is not publicly available and affordable price appears dubious. RNA biocontrols remain challenged by slower lethality and shorter residual activity than conventional insecticides. Sprayable dsRNA offers theoretical advantages over GM crops but requires deeper mechanistic understanding of RNAi pathways and environmental safety validation.

## 4. Conclusion

Although both CUAD biotechnology and double-stranded RNA technology utilize nucleic acids as active ingredients, they represent fundamentally distinct insecticidal approaches with divergent mechanisms and applications (Oberemok et al. 2018a). Oligonucleotide insecticides employ short, unmodified antisense DNA fragments delivered mainly via contact, operating through the DNA containment (DNAc) mechanism to degrade target ribosomal RNAs in pests. The deoxyribose backbone of DNA fragments confers greater resistance to environmental hydrolysis than RNA-based counterparts (Thorp, 2000), while their compact size (∼11 nucleotides) enables exceptional targeting precision. Ribosomal RNA – constituting approximately 80% of cellular RNA – serves as the optimal target due to its abundance and metabolic significance (Oberemok et al. 2019, 2023, 2025a; Gal’chinsky et al., 2024), with current efficacy being most pronounced against sap-feeding sternorrhynchans (Gal’chinsky et al. 2024; Oberemok et al. 2024a). In contrast, RNA biocontrols utilize longer double-stranded RNA molecules delivered through either contact or oral routes, functioning via the RNA interference (RNAi) pathway to degrade messenger RNAs. The ribose backbone of these molecules increases susceptibility to environmental hydrolysis (Williams et al. 2016), while their extended length (typically >200 bp) and subsequent enzymatic dicing into smaller fragments complicate selectivity control, necessitating careful risk assessment for non-target organisms (Roberts et al. 2015; Christiaens et al. 2022; Rank and Koch 2021; Liu et al. 2020). Presently, dsRNA insecticides demand optimized design algorithms and expanded target pest spectra, whereas CUADb-based oligonucleotide insecticides primarily require broader validation across additional pest groups (Oberemok et al. 2024d).

Both classes benefit from cost-effective synthesis: liquid-phase phosphoramidite methods for DNA oligonucleotides (Gal’chinsky et al. 2023; Oberemok et al. 2024b) and cell-free enzymatic production for dsRNA (Taning et al. 2020). Notably, while RNAi mechanisms were characterized years before practical dsRNA insecticides emerged, the DNAc mechanism was discovered following persistent applied research with antisense oligodeoxyribonucleotides in pest control. These complementary technologies will likely demonstrate optimal efficacy against specific pest taxa, offering extended utility through careful design and potential for synergistic integration in multi-target formulations. Their unprecedented safety profiles for non-target organisms and ecosystems – combining high selectivity, rapid environmental biodegradation, and innovative mechanisms – position nucleic acid-based insecticides as indispensable biomolecules for sustainable agriculture’s future.

## Author Contributions

Conceptualization, V.O.; software, V.O. and N.G.; validation, V.O., and N.G.; formal analysis, V.O. and N.G.; resources, V.O.; data curation, V.O., K.L., J.A., I.C., and N.G.; writing—original draft preparation, V.O. and N.G.; writing – review and editing, V.O., K.L., J.A., I.C., and N.G.; visualization, V.O. and N.G.; supervision, V.O. and N.G.; project administration, V.O.; funding acquisition, V.O.

## Institutional Review Board Statement

Not applicable.

## Informed Consent Statement

Not applicable.

## Data Availability Statement

No new data were created.

## Acknowledgments

We thank our many colleagues, too numerous to name, for the technical advances and lively discussions that prompted us to write this review. We apologize to the many colleagues whose work has not been cited. We are very much indebted to all anonymous reviewers and our colleagues from the lab on DNA technologies, PCR analysis, and creation of DNA insecticides (V.I. Vernadsky Crimean Federal University, Department of General Biology and Genetics), and OLINSCIDE BIOTECH LLC for valuable comments on our manuscript.

## References

Arora AK, Chung SH, Douglas AE (2021) Non-Target Effects of dsRNA Molecules in Hemipteran Insects. Genes. 12:407. 10.3390/genes12030407

Arya SK, Singh S, Upadhyay SK, Tiwari V, Saxena G, Verma PC (2021) RNAi□based gene silencing in Phenacoccus solenopsis and its validation by in planta expression of a double□stranded RNA. Pest Manag Sci 77(4):1796–1805. 10.1002/ps.6204

Bachellerie JP, Cavaille J, Huttenhofer A (2002) The expanding snoRNA world. Biochimie 84:775–790. 10.1016/S0300-9084(02)01402-5

Bachman P, Fischer J, Song Z, Urbanczyk-Wochniak E, Watson G (2020) Environmental Fate and Dissipation of Applied DsRNA in Soil, Aquatic Systems, and Plants. Front. Plant Sci 11:21. 10.3389/fpls.2020.00021

Barnes MA, Turner CR (2016) The ecology of environmental DNA and implications for conservation genetics. Conserv Genet 17:1–17. 10.1007/s10592-015-0775-4

Baum JA, Bogaert T, Clinton W, Heck GR, Feldmann P, Ilagan O, Johnson S, Plaetinck G, Munyikwa T, Pleau M, et al (2007) Control of coleopteran insect pests through RNA interference. Nat Biotechnol 25(11):1322–6. 10.1038/nbt1359

Bera P, Suby SB, Dixit S, Vijayan V, Kumar N, Sekhar JC, Vadassery J (2025) Identification of novel target genes for RNAi mediated management of the pest, Fall Armyworm (Spodoptera frugiperda, JE Smith). Crop Protection 187:106972. 10.1016/j.cropro.2024.106972

Bodemann RR, Rahfeld P, Stock M, Kunert M, Wielsch N, Groth M, Frick S, Boland W, Burse A (2012). Precise RNAi-mediated silencing of metabolically active proteins in the defence secretions of juvenile leaf beetles. Proceedings of the Biological Sciences. 279:4126–4134. 10.1098/rspb.2012.1342

BBolognesi R, Ramaseshadri P, Anderson J, Bachman P, Clinton W, Flannagan R, Ilagan O, Lawrence C, Levine S, Moar W, et al (2012) Characterizing the mechanism of action of double-stranded RNA activity against western corn rootworm (Diabrotica virgifera virgifera LeConte). PLoS One 7(10):e47534. 10.1371/journal.pone.0047534

Bramlett M, Plaetinck G, Maienfisch P (2020) RNA-based biocontrols—a new paradigm in crop protection. Engineering 6:522–527. 10.1007/10.1016/j.eng.2019.09.008

Brutscher LM, Daughenbaugh KF, Flenniken ML (2017) Virus and dsRNA-triggered transcriptional responses reveal key components of honey bee antiviral defense. Sci. Rep 7:1–15. 10.1038/s41598-017-06623-z

Buer B, DoCnitz J, Milner M, Mehlhorn S, Hinners C, Siemanowski-Hrach J, Ulrich JK, Grossmann D, Cedden D, Nauen R, et al (2024) Superior target genes and pathways for RNAi mediated pest control revealed by genome wide analysis in the red flour beetle Tribolium castaneum. bioRxiv. 10.1101/2024.01.24.577003

Cagliari D, Dias NP, Galdeano DM, dos Santos EÁ, Smagghe G, Zotti MJ (2019) Management of pest insects and plant diseases by non-transformative RNAi. Front Plant Sci 10. 10.3389/fpls.2019.01319

Camargo RA, Barbosa GO, Possignolo IP, Peres LE, Lam E, Lima JE (2016) Figueira, A.; Marques-Souza, H. RNA interference as a gene silencing tool to control Tuta absoluta in tomato (Solanum lycopersicum). PeerJ 4:e2673. 10.7717/peerj.2673

Cao M, Gatehouse, JA, Fitches EC (2018) A Systematic Study of RNAi Effects and dsRNA Stability in Tribolium castaneum and Acyrthosiphon pisum, Following Injection and Ingestion of Analogous dsRNAs. Int J Mol Sci 19:1079. 10.3390/ijms19041079

Carmell MA, Xuan Z, Zhang MQ, Hannon GJ (2002) The Argonaute family: tentacles that reach into RNAi, developmental control, stem cell maintenance, and tumorigenesis. Genes Dev 16:2733–2742. 10.1101/gad.1026102

Castellanos NL, Smagghe G, Sharma R, Oliveira EE, Christiaens O (2019) Liposome encapsulation and EDTA formulation of dsRNA targeting essential genes increase oral RNAi-caused mortality in the Neotropical stink bug Euschistus heros. Pest Manag Sci 75(2):537–548. 10.1002/ps.5167

Cedden D, Bucher G (2024) The quest for the best target genes for RNAi-mediated pest control. Insect Mol Biol 34(4):505–517. 10.1111/imb.12966

Cerio RJ, Vandergaast R, Friesen PD (2020) Host insect inhibitor-of-apoptosis SfIAP functionally replaces baculovirus IAP but is differentially regulated by its N-terminal leader. J Virol 84:11448–11460. 10.1128/jvi.01311-10

Chattopadhyay T, Biswal P, Lalruatfela A, Mallick B (2022) Emerging roles of PIWI-interacting RNAs (piRNAs) and PIWI proteins in head and neck cancer and their potential clinical implications Biochim. Biophys. Acta Rev Cancer 1877(5):188772. 10.1016/j.bbcan.2022.188772

Chen J, Peng Y, Zhang H, Wang K, Zhao C, Zhu G, Reddy Palli S, Han Z (2021) Off-target effects of RNAi correlate with the mismatch rate between dsRNA and non-target mRNA. RNA Biol 1–13. 10.1080/15476286.2020.1868680

Chen JZ, Jiang YX, Li MW, Li JW, Zha BH, Yang G (2021) Double-stranded RNA-degrading enzymes reduce the efficiency of RNA interference in Plutella xylostella. Insects 12(8):712. 10.3390/insects12080712

Christiaens O, Sweet J, Dzhambazova T, Urru I, Smagghe G, Kostov K, Arpaia S (2022) Implementation of RNAi-based arthropod pest control: environmental risks, potential for resistance and regulatory considerations. Journal of Pest Science 95:1–15. 10.1007/s10340-021-01439-3

Christiaens O, Swevers L, Smagghe G (2014) DsRNA Degradation in the Pea Aphid (Acyrthosiphon pisum) Associated with Lack of Response in RNAi Feeding and Injection Assay. Peptides 53:307–314. 10.1016/j.peptides.2013.12.014

Christiaens O, Whyard S, Vélez AM, Smagghe G (2020) Double-Stranded RNA Technology to Control Insect Pests: Current Status and Challenges. Front. Plant Sci 11:451. 10.3389/fpls.2020.00451

Cunningham DS, MacEachran D, Abshire JR, Dhamankar H, Iwuchukwu I, Gupta M, Moura ME, Sudharsan N, Skizim N, Jain R, et al (2020) Methods and compositions for nucleoside triphosphate and ribonucleic acid production. Washington, DC: U.S. Patent US20190144489A1.

Dalakouras A, Vlachostergios D (2021) Epigenetic approaches to crop breeding: current status and perspectives. J Exp Bot 72:5356–5371. 10.1093/jxb/erab227

Darlington M, Jurat-Fuentes JL, Kogel K, Rathore K, Smagghe G, Whyard S (2024) RNA interference in agriculture: methods, applications, and governance. CAST 72. 10.62300/IRNE9191

Deng ZL, MuCnch PC, Mreches R, McHardy AC (2022) Rapid and accurate identification of ribosomal RNA sequences via deep learning. Nucleic Acids Res 50(10):e60. 10.1093/nar/gkac112

Dias N, Stein CA (2002) Antisense oligonucleotides: basic concepts and mechanisms. Mol Cancer Ther 1(5):347–55.

Dietz-Pfeilstetter A, Mendelsohn M, Gathmann A, Klinkenbuß D (2021) Considerations and regulatory approaches in the USA and in the EU for dsRNA-based externally applied pesticides for plant protection. Front Plant Sci 12:974. 10.3389/fpls.2021.682387

Dubelman S, Fischer J, Zapata F, Huizinga K, Jiang C, Uffman J, Levine S, Carson D (2014) Environmental Fate of Double-Stranded RNA in Agricultural Soils. PLoS One. 9:e93155.

Environmental Protection Agency (EPA). Pesticide product label, SmartStax PRO Enlist Refuge https://www3.epa.gov/pesticides/chem_search/ppls/062719-00707-20170608.pdf. Accessed 19 April 2025

Evseev PV, Landysheva YG, Landyshev NN, Ignatov AN (2021) Presence of rRNA-like regions in Genbank viral sequences. 2021 IEEE Ural-Siberian Conference on Computational Technologies in Cognitive Science, Genomics and Biomedicine (CSGB) 310–314. 10.1109/CSGB53040.2021.9496035

Fire A, Xu S, Montgomery MK, Kostas SA, Driver SE, Mello CC (1998) Potent and specific genetic interference by double-stranded RNA in Caenorhabditis elegans. Nature 391:806–811.

Flenniken ML, Andino R (2013) Non-specific dsRNA-mediated antiviral response in the honey bee. PLoS One 8:e77263.

Gal’chinsky N, Useinov R, Yatskova E, Laikova K, Noviko I, Gorlov M, Trikoz N, Sharmagiy A, Plugatar Y, Oberemok V (2020) A breakthrough in the efficiency of contact DNA insecticides: rapid high mortality rates in the sap-sucking insects Dynaspidiotus britannicus Comstock and Unaspis euonymi Newstead. J Plant Prot Res 60(2):220–223. 10.24425/jppr.2020.133315

Gal’chinsky NV, Yatskova EV, Novikov IA, Sharmagiy AK, Plugatar YV, Oberemok VV (2024) Mixed insect pest populations of Diaspididae species under control of oligonucleotide insecticides: 3′-end nucleotide matters. Pesticide Biochemistry and Physiology 200:105838. 10.1016/j.pestbp.2024.105838

Gal’chinsky NV, Yatskova EV, Novikov IA, Useinov RZ, Kouakou NJ, Kouame KF, Kra KD, Sharmagiy AK, Plugatar YV, Laikova KV, et al (2023) Icerya purchasi Maskell (Hemiptera: Monophlebidae) Control Using Low Carbon Footprint Oligonucleotide Insecticides. Int J Mol Sci 24:11650. 10.3390/ijms241411650

Galli M, Feldmann F, Vogler UK, Kogel KH (2024) Can biocontrol be the game-changer in integrated pest management? A review of definitions, methods and strategies. J Plant Dis Prot 131:265–291. 10.1007/s41348-024-00878-1

Garbutt JS, Bellés X, Richards EH, Reynolds SE (2013) Persistence of double-stranded RNA in insect hemolymph as a potential determiner of RNA interference success: evidence from Manduca sexta and Blattella germanica. J Insect Physiol 59:171–8.

Gavrilova D, Grizanova E, Novikov I, Laikova E, Zenkova A, Oberemok V, Dubovskiy I (2025) Antisense DNA acaricide targeting pre-rRNA of two-spotted spider mite Tetranychus urticae as efficacy-enhancing agent of fungus Metarhizium robertsii. Journal of Invertebrate Pathology 211:108297. 10.1016/j.jip.2025.108297

GreenLight Biosciences. https://www.greenlightbiosciences.com/international-crop-network-validates-ledprona-as-a-new-mode-of-action-group/. Accessed 20 January 2025

Guan RB, Li HC, Fan YJ, Hu SR, Christiaens O, Smagghe G, Miao XX (2018) A Nuclease Specific to Lepidopteran Insects Suppresses RNAi. J Biol Chem 293:6011–6021. 10.1074/jbc.RA117.001553

Guo PP, Yang XB, Yang H, Zhou C, Long GY, Jin DC (2025) Knockdown of the β-N-acetylhexosaminidase genes by RNA interference inhibited the molting and increased the mortality of the white-backed planthopper, Sogatella furcifera. Pesticide Biochemistry and Physiology. 207:106216. 10.1016/j.pestbp.2024.106216

Guo W, Bai C, Wang Z, Wang P, Fan Q, Mi X, Wang L, He J, Pang J, Luo X, et al (2018) Double-stranded RNAs high-efficiently protect transgenic potato from Leptinotarsa decemlineata by disrupting juvenile hormone biosynthesis. J Agric Food Chem 66:11990–11999. 10.1021/acs.jafc.8b03914

Guo X, Wang Y, Sinakevitch I, Lei H, Smith BH (2018) Comparison of RNAi knockdown effect of tyramine receptor 1 induced by dsRNA and siRNA in brains of the honey bee, Apis mellifera. Journal of Insect Physiology 111:47–52. 10.1016/j.jinsphys.2018.10.005

Guo Y, Ji N, Bai L, Ma J, Li Z (2022) Aphid viruses: a brief view of a long history. Front Insect Sci 2:846716. 10.3389/finsc.2022.846716

Head GP, Carroll MW, Evans, SP, Rule DM, Willse AR, Clark TL, Storer NP, Flannagan RD, Samuel LW, Meinke LJ (2017) Evaluation of SmartStax and SmartStax PRO maize against western corn rootworm and northern corn rootworm: efficacy and resistance management. Pest Manag Sci 73:1883–1899. 10.1002/ps.4554

Hirai M, Terenius O, Li W, Faye I (2004) Baculovirus and dsRNA induce Hemolin, but no antibacterial activity, in Antheraea pernyi. Insect Mol Biol 13:399–405.

Hoang T, Foquet B, Rana S, Little DW, Woller DA, Sword GA, Song H (2022) Development of RNAi methods for the Mormon cricket, Anabrus simplex (Orthoptera: Tettigoniidae). Insects 13. 1010.3390/insects13080739

Irles P, Silva-Torres FA, Piulachs M.-D (2013) RNAi reveals the key role of Nervana 1 in cockroach oogenesis and embryo development. Insect Biochemistry and Molecular Biology 43:178–188. 10.1016/j.ibmb.2012.12.003

Ivashuta SI, Zhang Y, Wiggins EB, Ramaseshadri P, Segers GC, Johnson S, Meyer SE, Kerstetter RA, McNulty BC, Bolognesi R, et al (2015) Environmental RNAi in herbivorous insects. RNA 21:840–850. 10.1261/rna.048116.114

Jain RG, Robinson KE, Asgari S, Mitter N (2021) Current scenario of RNAi-based hemipteran control. Pest Manag Sci 77(5):2188–2196. 10.1002/ps.6153

Jain RG, Robinson KE, Fletcher SJ, Mitter N (2020) RNAi-Based Functional Genomics in Hemiptera. Insects 11(9):557. 10.3390/insects11090557

Khajuria C, Ivashuta S, Wiggins E, Flagel L, Moar W, Pleau M, Miller K, Zhang Y, Ramaseshadri P, Jiang C et al (2018) Development and characterization of the first dsRNA-resistant insect population from western corn rootworm, Diabrotica virgifera virgifera LeConte. PLoS One 13(5):e0197059. 10.1371/journal.pone.0197059

Koeppe S, Kawchuk L, Kalischuk M (2023) RNA Interference Past and Future Applications in Plants. Int J Mol Sci 24:9755. 10.3390/ijms24119755

Koo J, Palli SR (2024) Recent advances in understanding of the mechanisms of RNA interference in insects. Insect Molecular Biology 1–14. 10.1111/imb.12941

Kumar H, Gal’chinsky N, Sweta V, Negi N, Filatov R, Chandel A, Ali J, Oberemok V, Laikova K (2025) Perspectives of RNAi, CUADb and CRISPR/Cas as Innovative Antisense Technologies for Insect Pest Control: From Discovery to Practice. Insects 16:746. 10.3390/insects16070746

Kumar H, Sharma M, Chandel A (2022) DNA Insecticides: Future of Crop Protection. Agriculture & Food: e-Newsletter 4(6):551.

Li L, Jing A, Xie M, Li S, Ren C (2021) Applications of RNA interference in American cockroach. Journal of Visualized Experiments (178):e63380. 10.3791/63380

Li N, Xu X, Li J, Hull JJ, Chen L, Liang G (2024) A spray-induced gene silencing strategy for Spodoptera frugiperda oviposition inhibition using nanomaterial-encapsulated dsEcR. International Journal of Biological Macromolecules 281:136503

Liu S, Jaouannet M, Dempsey DA, Imani J, Coustau C, Kogel KH (2020) RNA-based technologies for insect control in plant production. Biotechnol Adv 39:107463. 10.1016/j.biotechadv.2019.107463

Lomate PR, Bonning BC (2016) Distinct Properties of Proteases and Nucleases in the Gut, Salivary Gland and Saliva of Southern Green Stink Bug, Nezara viridula. Sci Rep 6:27587. 10.1038/srep27587

Lu J, Shen J (2024) Target genes for RNAi in pest control: A comprehensive overview. Entomologia Generalis 44(1):95–114. 10.1127/entomologia/2023/2207

Mao J, Zeng F (2014) Plant-mediated RNAi of a gap gene-enhanced tobacco tolerance against the Myzus persicae. Transgenic Res 23:145–152. 10.1007/s11248-013-9739-y

Martinez Z, De Schutter K, Van Damme EJM, Vokgel E, Wynant N, Vanden Broeck J, Christiaens O, Smagghe G (2021) Accelerated delivery of dsRNA in lepidopteran midgut cells by a Galanthus nivalis lectin (GNA)-dsRNA-binding domain fusion protein. Pestic Biochem Phys 175:104853. 10.1016/j.pestbp.2021.104853

Miller SC, Brown SJ, Tomoyasu Y (2008) Larval RNAi in Drosophila? Development Genes and Evolution 218:505–510. 10.1007/s00427-008-0238-8

Moar W, Khajuria C, Pleau M, Ilagan O, Chen M, Jiang C, Price P, McNulty B, Clark T, Head G (2017) Cry3Bb1-Resistant Western Corn Rootworm, Diabrotica virgifera virgifera (LeConte) Does Not Exhibit Cross-Resistance to DvSnf7 dsRNA. PLoS One 12(1):e0169175. 10.1371/journal.pone.0169175

Mogilicherla K, Roy A (2023) Epigenetic regulations as drivers of insecticide resistance and resilience to climate change in arthropod pests. Front Genet 13:1044980. 10.3389/fgene.2022.1044980

Müller M, Fazi F, Ciaudo C (2020) Argonaute Proteins: From Structure to Function in Development and Pathological Cell Fate Determination. Front Cell Dev Biol 7:360. 10.3389/fcell.2019.00360.

Narva K, Toprak U, Alyokhin A, Groves R, Jurat-Fuentes JL, Moar W, Nauen R, Whipple S, Head G (2025) Insecticide resistance management scenarios differ for RNA-based sprays and traits. Insect Mol Biol 34(4):518–526. 10.1111/imb.12986

Nitnavare RB, Bhattacharya J, Singh S, Kour A, Hawkesford MJ, Arora N (2021) Next Generation dsRNA-Based Insect Control: Success So Far and Challenges. Front Plant Sci 12:673576. 10.3389/fpls.2021.673576

Novikov A, Yatskova E, Bilyk A, Puzanova Y, Sharmagiy A, Oberemok V (2023) Efficient Control of the Obscure Mealybug Pseudococcus viburni with DNA Insecticides. In Vitro Cellular & Developmental Biology-Animal 59:92–108. 10.1007/s11626-023-00795-x

Nyadar PM, Oberemok V, Omelchenko A, Kerimova S, Seidosmanova E, Krasnodubiets A, Shumskykh M, Bekirova V, Galchinsky N, Vvdensky V (2019) DNA Insecticides: The Effect of Concentration on Non-Target Plant Organisms Such as Wheat (Triticum aestivum L.). J Plant Prot Res 59:60–68. 10.24425/jppr.2019.126038

Obbard DJ, Gordon KH, Buck AH, Jiggins FM (2009) The evolution of RNAi as a defence against viruses and transposable elements. Philos Trans R Soc Lond B Biol Sci 364(1513):99–115. 10.1098/rstb.2008.0168

Oberemok V, Gal’chinsky N, Novikov I, Sharmagiy A, Yatskova E, Laikova E, Plugatar Y (2025a) Ribosomal RNA-Specific Antisense DNA and Double-Stranded DNA Trigger rRNA Biogenesis and Insecticidal Effects on the Insect Pest Coccus hesperidum. Int J Mol Sci 26:7530. 10.3390/ijms26157530

Oberemok V, Laikova K, Andreeva O, Dmitrienko A, Rybareva T, Ali J, Gal’chinsky N (2025c) DNA-Programmable Oligonucleotide Insecticide Eriola-11 Targets Mitochondrial 16S rRNA and Exhibits Strong Insecticidal Activity Against Woolly Apple Aphid (Eriosoma lanigerum) Hausmann. Int J Mol Sci 26:7486. 10.3390/ijms26157486

Oberemok VV (2008) Method of elimination of phyllophagousinsects from order Lepidoptera. UA Patent 36445.

Oberemok VV, Gal’chinsky NV, Useinov RZ, Novikov IA, Puzanova YV, Filatov RI, Kouakou NJ, Kouame KF, Kra KD, Laikova KV, (2023) Four Most Pathogenic Superfamilies of Insect Pests of Suborder Sternorrhyncha: Invisible Superplunderers of Plant Vitality. Insects 14(5):462. 10.3390/insects14050462

Oberemok VV, Laikova K, Shumskykh M, Kenyo I, Kasich I, Deri K, Seidosmanova E, Krasnodubets K, Bekirova V, Gal’chinsky N (2018) A primary attempt of Leptinotarsa decemlineata control using contact DNA insecticide based on short antisense oligonucleotide of its CYP6B gene. J Plant Prot Res 58:106–108. 10.24425/119124

Oberemok VV, Laikova KV, Andreeva OA, Gal’chinsky NV (2024) Biodegradation of insecticides: oligonucleotide insecticides and double-stranded RNA biocontrols paving the way for eco-innovation. Front Environ Sci 12. 10.3389/fenvs.2024.1430170

Oberemok VV, Laikova KV, Gal’chinsky NV (2024) Contact unmodified antisense DNA (CUAD) biotechnology: list of pest species successfully targeted by oligonucleotide insecticides. Front Agron 6:1415314. 10.3389/fagro.2024.1415314

Oberemok VV, Laikova KV, Gal’chinsky NV, Useinov RZ, Novikov IA, Temirova ZZ, Shumskykh MN, Krasnodubets AM, Repetskaya AI, Dyadichev VV, et al (2019) DNA insecticide developed from the Lymantria dispar 5.8S ribosomal RNA gene provides a novel biotechnology for plant protection. Sci Rep 9(1):6197. 10.1038/s41598-019-42688-8

Oberemok VV, Laikova KV, Repetskaya AI, Kenyo IM, Gorlov MV, Kasich IN, Krasnodubets AM, Gal’chinsky NV, Fomochkina II, Zaitsev AS, et al (2018) A Half-Century History of Applications of Antisense Oligonucleotides in Medicine, Agriculture and Forestry: We Should Continue the Journey. Molecules 23(6):1302. 10.3390/molecules23061302

Oberemok VV, Laikova KV, Zaitsev AS, Shumskykh MN, Kasich IN, Gal’chinsky NV, Bekirova VV, Makarov VV, Agranovsky AA, Gushchin VA et al (2017) Molecular Alliance of Lymantria dispar Multiple Nucleopolyhedrovirus and a Short Unmodified Antisense Oligonucleotide of Its Anti-Apoptotic IAP-3 Gene: A Novel Approach for Gypsy Moth Control. Int J Mol Sci 18:2446. 10.3390/ijms18112446

Oberemok VV, Laikova VK, Zaitsev SA, Nyadar MP, Shumskykh NM, Gninenko IY (2015) DNA insecticides based on iap3 gene fragments of cabbage looper and gypsy moth nuclear polyhedrosis viruses show selectivity for non-target insects. Arch Biol Sci 67:785–792.

Oberemok VV, Novikov IA, Yatskova EV, Bilyk AI, Sharmagiy AK, Gal’chinsky NV (2024c) Potent and selective ‘genetic zipper’ method for DNA-programmable plant protection: innovative oligonucleotide insecticides against Trioza alacris Flor. Chem Biol Technol Agric 11:144. 10.1186/s40538-024-00668-9

Oberemok VV, Nyadar P, Zaytsev O, Levchenko N, Shiyntum H, Omelchenko O (2013) Pioneer evaluation of the possible side effects of the DNA insecticides on wheat (Triticum aestivum L.). Int J Biochem Biophys 1:57–63.

Oberemok VV, Puzanova YV, Gal’chinsky NV (2024b) The ‘genetic zipper’ method offers a cost-effective solution for aphid control. Front Insect Sci 4:1467221. 10.3389/finsc.2024.1467221

Oberemok VV, Useinov RZ, Skorokhod OA, Gal’chinsky NV, Novikov IA, Makalish TP, Yatskova EV, Sharmagiy AK, Golovkin IO, Gninenko YI, et al (2022) Oligonucleotide Insecticides for Green Agriculture: Regulatory Role of Contact DNA in Plant–Insect Interactions. Int J Mol Sci 23:15681. 10.3390/ijms232415681

OECD. Considerations for the Environmental Risk Assessment of the Application of Sprayed or Externally Applied dsRNA-Based Pesticides. Series on Pesticides No. 104, ENV/JM/MONO 2020. Paris: OECD (2020) p. 26. https://www.oecd.org/content/dam/oecd/en/publications/reports/2020/09/considerations-for-the-environmental-risk-assessment-of-the-application-of-sprayed-or-externally-applied-ds-rna-based-pesticides_898e0d9f/576d9ebb-en.pdf. Accessed 12 August 2025

Palli SR (2023) RNAi turns 25: contributions and challenges in insect science. Front Insect Sci 3:1209478. 10.3389/finsc.2023.1209478

Pallis S, Alyokhin A, Manley B, Rodrigues T, Barnes E, Narva K (2023) Effects of Low Doses of a Novel dsRNA-based Biopesticide (Calantha) on the Colorado Potato Beetle. J Econ Entomol 116(2):456–461. 10.1093/jee/toad034

Parker KM, Barragán Borrero V, van Leeuwen DM, Lever MA, Mateescu B, Sander M (2019) Environmental Fate of RNA Interference Pesticides: Adsorption and Degradation of Double-Stranded RNA Molecules in Agricultural Soils. Environ Sci Technol 53:3027–3036.

Peng Y, Wang K, Fu W, Sheng C, Han Z (2018) Biochemical Comparison of dsRNA Degrading Nucleases in Four Different Insects. Front Physiol 9:624. 10.3389/fphys.2018.00624

Powell M, Pyati P, Cao M, Bell H, Gatehouse JA, Fitches E (2017) Insecticidal effects of dsRNA targeting the Diap1 gene in dipteran pests. Sci Rep 7:1–13.

Powell ME, Bradish HM, Gatehouse JA, Fitches EC (2017) Systemic RNAi in the small hive beetle Aethina tumida Murray (Coleoptera: Nitidulidae), a serious pest of the European honey bee Apis mellifera. Pest Manag Sci 73:53–63. 10.1002/ps.4365

Prentice K, Christiaens O, Pertry I, Bailey A, Niblett C, Ghislain M, Gheysen G, Smagghe G (2017) RNAi□based gene silencing through dsRNA injection or ingestion against the African sweet potato weevil Cylas puncticollis (Coleoptera: Brentidae). Pest Manag Sci 73(1):44–52.

Puzanova YV, Novikov IA, Bilyk AI, Sharmagiy AK, Plugatar YV, Oberemok VV (2023) Perfect Complementarity Mechanism for Aphid Control: Oligonucleotide Insecticide Macsan-11 Selectively Causes High Mortality Rate for Macrosiphoniella sanborni Gillette. Int J Mol Sci 24:11690. 10.3390/ijms241411690

Ramaseshadri P, Segers G, Flannagan R, Wiggins E, Clinton W, Ilagan O, McNulty B, Clark T, Bolognesi R (2013) Physiological and cellular responses caused by RNAi-mediated suppression of Snf7 orthologue in western corn rootworm (Diabrotica virgifera virgifera) larvae. PLoS One 8(1):e54270. 10.1371/journal.pone.0054270

Rank AP, Koch A (2021) Lab-to-Field Transition of RNA Spray Applications - How Far Are We? Front Plant Sci 12:755203. 10.3389/fpls.2021.755203

Rawlinson SM, Zhao T, Rozario AM, Rootes CL, McMillan PJ, Purcell AW, Woon A, Marsh GA, Lieu KG, Wang LF, et al (2018) Viral regulation of host cell biology by hijacking of the nucleolar DNA-damage response. Nat Commun 9(1):3057. 10.1038/s41467-018-05354-7

Reinders JD, Moar WJ, Head GP, Hassan S, Meinke LJ (2023) Effects of SmartStax® and SmartStax® PRO maize on western corn rootworm (Diabrotica virgifera virgifera LeConte) larval feeding injury and adult life history parameters. PLoS One. 18(7):e0288372. 10.1371/journal.pone.0288372

Roberts AF, Devos, Y, Lemgo GNY, Zhou X (2015) Biosafety research for non-target organism risk assessment of RNAi-based GE plants. Front Plant Sci 6:958. 10.3389/fpls.2015.00958

Rodrigues T, Sridharan K, Manley B, Cunningham D, Narva K (2021) Development of dsRNA as a Sustainable Bioinsecticide: From Laboratory to Field. In B. Rauzan and B. A. Lorsbach (eds.). Crop Protection Products for Sustainable Agriculture. pp. 65–82. 10.1021/bk-2021-1390.ch005

Rodrigues TB, Dhandapani RK, Duan JJ, Palli SR (2017) RNA interference in the Asian Longhorned beetle: identification of key RNAi genes and reference genes for RT-qPCR. Scientific Reports 7:8913. 10.1038/s41598-017-08813-1

Rodrigues TB, Mishra SK, Sridharan K, Barnes ER, Alyokhin A, Tuttle R, Kokulapalan W, Garby D, Skizim N, Tang YW, et al (2021) First sprayable double-stranded RNA-bases biopesticide product target Type-5 Colorado potato beetle. Front Plant Sci 12:728652. 10.3389/fpls.2021.728652

Rodrigues TB, Rieske LK, Duan J, Mogilicherla K, Palli SR (2017) Development of RNAi method for screening candidate genes to control emerald ash borer, Agrilus planipennis. Scientific Reports 7:7379. 10.1038/s41598-017-07605-x

Salvetti A, Greco A (2024) Viruses and the nucleolus: the fatal attraction. Biochim Biophys Acta 1842(6):840–7. 10.1016/j.bbadis.2013.12.010

Santos D, Vanden Broeck J, Wynant N (2014) Systemic RNA interference in locusts: reverse genetics and possibilities for locust pest control. Current Opinion in Insect Science 6:9–14. 10.1016/j.cois.2014.09.013

Schellens S, Lenaerts C, Pérez Baca MDR, Cools D, Peeters P, Marchal E, Vanden Broeck J (2022) Knockdown of the Halloween genes Spook, Shadow and Shade influences oocyte development, egg shape, oviposition and hatching in the desert locust. Int J Mol Sci 23(16):9232. 10.3390/ijms23169232

Sharma VK, Sharma RK, Singh SK (2014) Antisense oligonucleotides: modifications and clinical trials. Med Chem Comm 5:1454–1471.

Shen GM, Song CG, Ao YQY, Xiao YH, Zhang YJ, Pan Y, He L (2017) Transgenic cotton expressing CYP392A4 double-stranded RNA decreases the reproductive ability of Tetranychus cinnabarinus. Insect Sci 24:559–568. 10.1111/1744-7917.12346

Sparks TC, Storer N, Porter A, Slater R, Nauen R (2021) Insecticide resistance management and industry: the origins and evolution of the Insecticide Resistance Action Committee (IRAC) and the mode of action classification scheme. Pest Manag Sci 77(6):2609–2619. 10.1002/ps.6254

Sun Z, Liu J, Chen Y, Zhang J, Zhong G (2023) RNAi-mediated knockdown of α-Spectrin depresses reproductive performance in female Bactrocera dorsalis. Pesticide Biochemistry and Physiology 196:105611.

Svoboda P (2020) Key Mechanistic Principles and Considerations Concerning RNA Interference. Front Plant Sci 11:1237. 10.3389/fpls.2020.01237

Szaflarski W, Leśniczak-Staszak M, Sowiński M, Ojha S, Aulas A, Dave D, Malla S, Anderson P, Ivanov P, Lyons SM (2022) Early rRNA processing is a stress-dependent regulatory event whose inhibition maintains nucleolar integrity. Nucleic Acids Res 50(2):1033–1051. 10.1093/nar/gkab1231

Taning CNT, Arpaia S, Christiaens O, Dietz-Pfeilstetter A, Jones H, Mezzetti B, Sabbadini S, Sorteberg H, Sweet J, Ventura V, et al (2020) RNA-Based Biocontrol Compounds: Current Status and Perspectives to Reach the Market. Pest Manag Sci 76:841–845. 10.1002/ps.5686

Terenius O, Papanicolaou A, Garbutt JS, Eleftherianos I, Huvenne H, Kanginakudru S, Albrechtsen M, An C, Aymeric JL, Barthel A, et al (2011) RNA interference in Lepidoptera: an overview of successful and unsuccessful studies and implications for experimental design. J Insect Physiol 57:231–245

Thorp HH (2000) The importance of being r: greater oxidative stability of RNA compared with DNA. Chem Biol 2000 7(2):33–6. 10.1016/s1074-5521(00)00080-6

Tomoyasu Y, Miller C, Tomita S, Schoppmeier M, Grossmann D, Bucher G (2008) Exploring systemic RNA interference in insects: a genomewide survey for RNAi genes in Tribolium. Genome biology 9(1).

TriLink BioTechnologies (2025) Feasibility of Antisense Oligonucleotides as DNAInsecticides. https://www.trilinkbiotech.com/blog/feasibility-of-antisense-oligonucleotides-as-dna-insecticides/ Accessed 19 April 2025

Ulrich J, Dao VA, Majumdar U, Schmitt-Engel C, Schwirz J, Schultheis D, StroChlein N, Troelenberg N, Grossmann D, Richter T, et al (2015) Large scale RNAi screen in Tribolium reveals novel target genes for pest control and the proteasome as prime target. BMC Genomics 16:674.

Useinov RZ, Gal’chinsky N, Yatskova E, Novikov I, Puzanova Y, Trikoz N, Sharmagiy A, Plugatar Y, Laikova K, Oberemok V (2020) To bee or not to bee: creating DNA insecticides to replace non-selective organophosphate insecticides for use against the soft scale insect Ceroplastes japonicus Green. J Plant Prot Res 60:406–409.

Vaccari T, Rusten TE, Menut L, Nezis IP, Brech A, Stenmark H, Bilder D (2009) Comparative analysis of ESCRT-I, ESCRT-II and ESCRT-III function in Drosophila by efficient isolation of ESCRT mutants. J Cell Sci 122(Pt 14):2413–23. 10.1242/jcs.046391

van den Berg H, daSilva Bezerra HS, Al-Eryani S, Chanda E, Nagpal BN, Knox TB, Velayudhan R, Yadav RS (2021) Recent trends in global insecticide use for disease vector control and potential implications for resistance management. Sci Rep 11(1):23867. 10.1038/s41598-021-03367-9

Vilkhovoy M, Adhikari A, Vadhin S, Varner JD (2020) The Evolution of Cell Free Biomanufacturing. Processes 8:675. 10.3390/pr8060675

Wang W, Han BW, Tipping C, Ge DT, Zhang Z, Weng Z, Zamore PD (2015) Slicing and binding by Ago3 or Aub trigger Piwi-bound piRNA production by distinct mechanisms. Mol Cell 59:819–30

Wang XW, Blanc S (2021) Insect transmission of plant single-stranded DNA viruses. Annu Rev Entomol 66:389–405. 10.1146/annurev-ento-060920-094531

Wang Y, Zhang H, Li H., Miao X (2011) Second-generation sequencing supply an effective way to screen RNAi targets in large scale for potential application in pest insect control. PloS One 6:e18644. 10.1371/journal.pone.0018644

Whalon ME, Mota-Sanchez RM, Hollingworth RM (2015) Arthropods Resistant to Pesticides Database (ARPD). http://www.pesticideresistance.org. Accessed 8 December 2015

Will CL, Luhrmann R (2001) Spliceosomal UsnRNP biogenesis, structure and function. Curr Opin Cell Biol 13:290–301. 10.1016/S0955-0674(00)00211-8

Williams JS, Lujan SA, Kunkel TA (2016) Processing ribonucleotides incorporated during eukaryotic DNA replication. Nat Rev Mol Cell Biol 17(6):350–63. 10.1038/nrm.2016.37

Wu J, Yang J, Cho WC, Zheng Y (2020) Argonaute proteins: Structural features, functions and emerging roles. J Adv Res 24:317–324. 10.1016/j.jare.2020.04.017.

Wu W, Shan HW, Li JM, Zhang CX, Chen JP, Mao Q (2022) Roles of bacterial symbionts in transmission of plant virus by hemipteran vectors. Front Microbiol 13:805352. 10.3389/fmicb.2022.805352

Xiong Y, Zeng H, Zhang Y, Xu D, Qiu D (2013) Silencing the HaHR3 gene by transgenic plant-mediated RNAi to disrupt Helicoverpa armigera development. Int J Biol Sci 9:370–381. 10.7150/ijbs.5929

Yoshiyama N, Tojo K, Hatakeyama M (2013) A survey of the effectiveness of non-cell autonomous RNAi throughout development in the sawfly, Athalia rosae (Hymenoptera). Journal of Insect Physiology 59:400–407. 10.1016/j.jinsphys.2013.01.009

Zaitsev AS, Omel’chenko OV, Nyadar PM, Oberemok VV (2015) Influence of DNA oligonucleotides used as insecticides on biochemical parameters of Quercus robur and Malus domestica. Bull Transylvania Univ Bras Ser II For Wood Ind Agric Food Eng 8:37–46.

Zha W, Peng X, Chen R, Du B, Zhu L, He G (2011) Knockdown of midgut genes by dsRNA-transgenic plant-mediated RNA interference in the hemipteran insect Nilaparvata lugens. PloS One 6(5):e20504. 10.1371/journal.pone.0020504

Zheng J, Xu Y (2023) A Review: Development of Plant Protection Methods and Advances in Pesticide Application Technology in Agro-Forestry Production. Agriculture 13:2165. 10.3390/agriculture13112165

Zotti M, dos Santos EA, Cagliari D, Christiaens O, Taning CNT, Smagghe G (2018) RNA interference technology in crop protection against arthropod pests, pathogens and nematodes. Pest Manag Sci 74:1239–1250.

